# Speech-evoked brain activity is more robust to competing speech when it is spoken by someone familiar

**DOI:** 10.1101/2020.03.03.975409

**Authors:** Emma Holmes, Ingrid S. Johnsrude

**Affiliations:** The Brain and Mind Institute, University of Western Ontario, London, Ontario, N6A 3K7, Canada; School of Communication Sciences and Disorders, University of Western Ontario, London, Ontario, London, N6G 1H1, Canada

**Keywords:** Speech, voice, familiarity, attention, auditory cortex, fMRI

## Abstract

People are much better at understanding speech when it is spoken by a familiar talker—such as a friend or partner—than when the interlocutor is unfamiliar. This provides an opportunity to examine the substrates of intelligibility and familiarity, independent of acoustics. Is the familiarity effect evident as early as primary auditory cortex, or only at later processing stages? Here, we presented sentences spoken by naturally familiar talkers (the participant’s friend or partner) and unfamiliar talkers (the friends or partners of other participants). We compared multivariate activity in speech-sensitive regions of cortex between conditions in which target sentences were presented alone and conditions in which the same target sentences were presented at the same time as a competing sentence. Using representational similarity analysis (RSA), we demonstrate that the pattern of activity evoked by a spoken sentence is less degraded by the presence of a competing sentence when it is spoken by a friend or partner than by someone unfamiliar; the results cannot be explained by acoustic differences since familiar and unfamiliar talkers were nearly identical across the group. This familiar-voice advantage is most prominent in nonprimary auditory cortical areas, along the posterior superior and middle temporal gyri. Across participants, the magnitude of the familiar-unfamiliar RSA difference correlates with the familiar-voice benefit to intelligibility. Overall, our results demonstrate that experience-driven improvements in intelligibility are associated with enhanced patterns of neural activity in nonprimary auditory cortical areas.

**Significance statement:** Speech is a complex signal, and we do not yet fully understand how the content of a spoken sentence is encoded in cortex. Here, we used a novel approach based on analysing multivariate activity: we compared activity evoked by highly intelligible sentences presented alone and by the same sentences presented with a competing masker. The distributed pattern of activity in speech-sensitive regions of the brain was more similar between the alone and masker conditions when the target sentence was spoken by someone familiar—the participant’s friend or partner—than someone unfamiliar. This metric correlated with the intelligibility of the familiar voice. These results imply that the spatial pattern of activity in speech-sensitive regions reflects the intelligibility of a spoken sentence.

## Introduction

In everyday life, speech can be difficult to understand when other conversations take place at the same time. Being familiar with a conversational partner is associated with better speech intelligibility when a competing talker is present (1–8). This familiar-voice benefit is substantial—participants report 10–20% more sentences correctly when they are spoken by their friend or spouse than when they are spoken by someone they have never met, and this cannot be explained by different acoustics of familiar and unfamiliar voices, since in a subset of these studies, familiar and unfamiliar voices are identical over the group (1–4). Despite this large and consistent benefit to speech intelligibility, the neural mechanisms by which familiarity improves intelligibility are currently unknown.

Previous functional imaging studies have typically manipulated intelligibility by changing speech acoustics or lexical predictability. Studies manipulating speech acoustics have demonstrated that better speech intelligibility is associated with greater activity in the superior temporal sulcus (STS; 9–11) and superior temporal gyrus (STG; 12, 13). However, in these studies, it is difficult to disentangle effects of acoustics from differences in intelligibility. A study manipulating lexical predictability (14) measured responses to degraded speech when it was preceded by a visual word prime: speech was rated as clearer when the word prime matched the spoken word than when it was different. The improvement in speech clarity for speech preceded by matching word primes was associated with greater activity in bilateral STS and left STG, including cytoarchitectonically defined primary auditory cortex. Overall, these findings are consistent with the idea that more intelligible speech is associated with greater activity along the superior temporal lobe, including primary auditory cortex.

Recent neuroimaging analyses have moved beyond simple activation maps to characterise the overall pattern of activity within a brain area. Whereas traditional (univariate) approaches to fMRI analyses spatially smooth the data and compare average activity across areas, comparing unsmoothed activity using multivariate rather than univariate analyses improves sensitivity to distributed activity (15, 16). One popular approach is Representational Similarity Analysis (RSA; 17, 18), which quantifies the difference between conditions as the ‘distance’ in representational space between their associated multivariate activities. One advantage of these approaches is that they can detect between-condition differences in the pattern of activity across voxels, even when average activity is the same. This approach is well-suited to study speech intelligibility—to quantify the difference in distributed activity between clear and degraded speech. Given that familiarity with a talker improves intelligibility in noise—in other words, making the intelligibility of speech-in-noise more similar to that of speech-in-quiet—we hypothesised that we could distinguish areas sensitive to intelligibility from those sensitive to acoustics by comparing activation patterns for familiar compared to unfamiliar voices, when the voices themselves are as identical as possible across the group. We conclude that regions exhibiting multivariate activity that is more similar for speech-in-noise and the same speech presented alone when the talker is familiar, compared to unfamiliar, are sensitive specifically to intelligibility of speech. We can use this marker of intelligibility sensitivity to functionally parcellate auditory cortex, using it to distinguish between regions that reflect experience-dependent enhancement of intelligibility, from those that do not.

We used high-resolution fMRI, combined with RSA, to measure activity that was elicited by sentences that were presented alone and by the same sentences that were presented simultaneously with a competing sentence spoken by a different talker. Comparing the multivariate activity in these two conditions revealed the extent to which the pattern of brain activity was disrupted by a competing (unfamiliar) talker. We compared conditions in which participants listened to speech spoken by a familiar talker (their friend or partner), and by unfamiliar takers, who were the friends and partners of other participants. Thus, familiar and unfamiliar stimuli were identical across the group.

## Results

### Replication of familiar-voice benefit to intelligibility

In the scanner, familiar and unfamiliar voices were either presented alone (“Familiar Alone” and “Unfamiliar Alone” conditions) or simultaneously with a competing sentence spoken by a different talker who was always unfamiliar (“Familiar Masked” and “Unfamiliar Masked” conditions). After the sentences had ended, participants saw a probe sentence presented visually on the screen and were asked to report whether the target sentence was the same as or different than the probe (see Figure 1).

**Figure 1.**
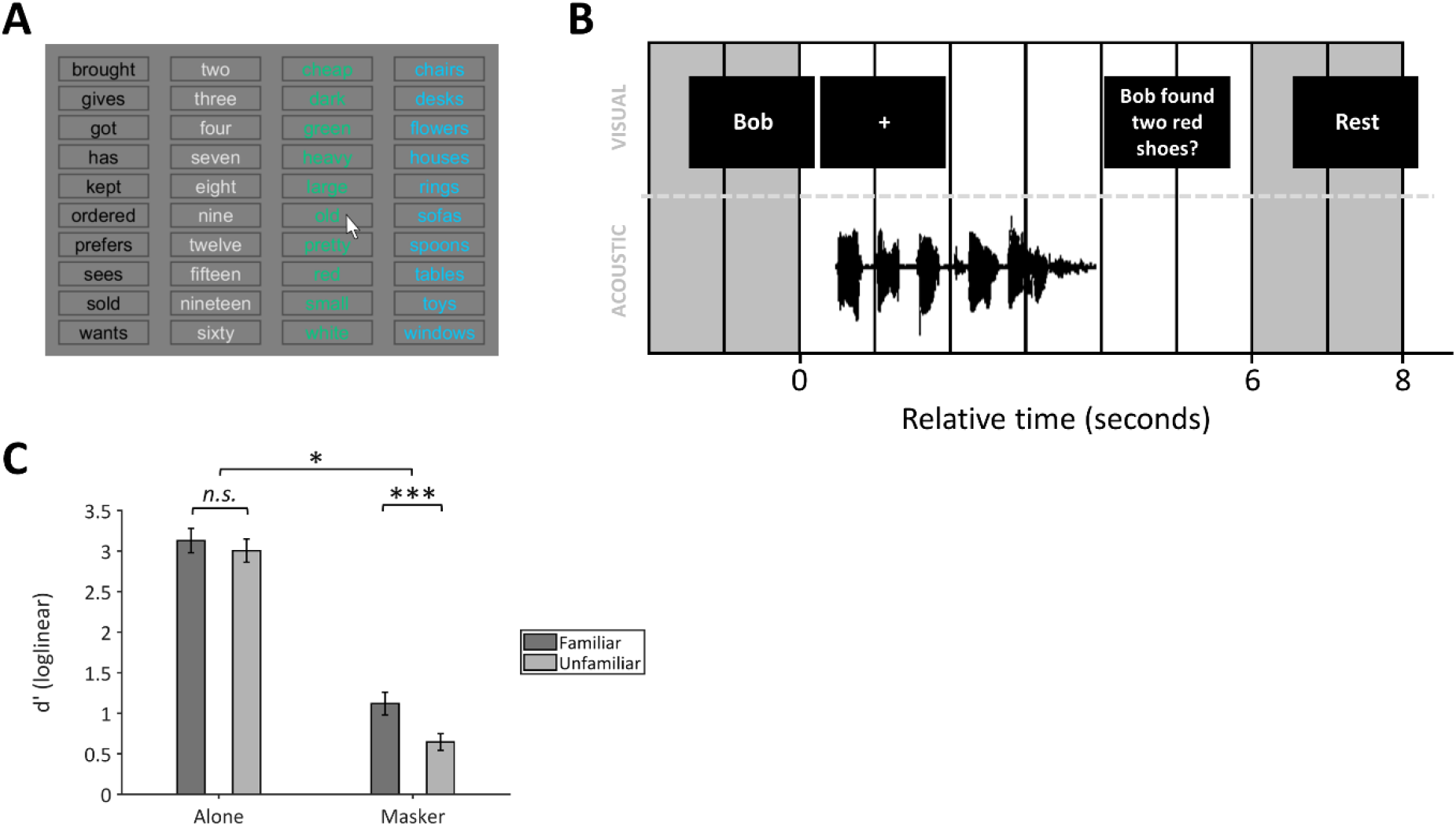
Overview of design. (A) Schematic of the response screen used for the tasks conducted outside the scanner (i.e., pre-scan and post-scan behavioural tasks). On each trial, participants clicked one word from each column, in any order. (B) Schematic of trial structure for the functional runs of the MRI session. An example trial is displayed, with the visual stimuli on the upper row, and the acoustic stimulus on the lower row. Each grey bar indicates one fMRI volume acquisition. White bars indicate ‘silent’ scans without volume acquisition. The acoustic stimulus is always presented during the ‘silent’ scans. The cue (“Bob”) for the example trial is presented during the volume acquisitions for the previous scan, and the cue (“Rest”) for the next (Silence) trial is presented during the volume acquisitions at the end of the trial. (C) Behavioural sensitivity (d′ with loglinear correction; *N* = 27) during the functional runs of the MRI session. Error bars display ±1 standard error of the mean. [*** *p* < 0.001; ** *p* < 0.01; * *p* < 0.05; *n.s.* not significant]

Participants performed well in the Familiar Alone (mean = 92.5%, S.E. = 1.9) and Unfamiliar Alone (mean = 91.4%, S.E. = 2.0) conditions. They performed lower in the Familiar Masked condition (mean = 69.5%, S.E. = 2.4) and worst in the Unfamiliar Masked condition (mean = 61.0%, S.E. = 1.9).

We replicated the behavioural familiar-voice intelligibility benefit in the MRI session using our new task (Figure 1C), as confirmed by the results of a 2 × 2 ANOVA (factors: Familiarity and Masker). As expected, the main effect of Familiarity [F(1, 26) = 16.29, *p* < .001, *ω*_*p*_^2^ = .35], the main effect of Masker [F(1, 26) = 270.60, p < .001, *ω*_*p*_^2^ = .91], and the Familiarity-Masker interaction [F(1, 26) = 6.99, *p* = .014, *ω*_*p*_^2^ = .18] were all significant. Paired-samples t-tests showed better d′ in the Familiar Masked than the Unfamiliar Masked condition [*t*(26) = 4.12, *p* < .001, *d*_*z*_ = .79], but no difference between the Familiar Alone and Unfamiliar Alone conditions [*t*(26) = 1.53, *p* = .14, *d*_*z*_ = .29].

We also tested intelligibility of the Familiar Masked and Unfamiliar Masked materials after the scan: Participants were asked to click words on the screen corresponding to the words they heard in the target sentence (Figure 1A). Across participants, the percentage of words reported correctly after the scanning session correlated with d′ in the scanner, both for the Familiar Masked [*r* = .68, *p* < .001; 95% CI = .39–.84] and Unfamiliar Masked [*r* = .58, *p* = .001; 95% CI = .26–.79] conditions.

### Pattern of activity is more robust for familiar voices

We used RSA to test the dissimilarity of multivariate activity between the target sentence in the Masked conditions and the same sentences presented in the Alone conditions. This analysis tested the dissimilarity across all of the voxels in a ‘Speech Perception’ region of interest (ROI), which was extracted from the Neurosynth database. The correlation distance was significantly smaller between the Familiar Masked and Familiar Alone conditions (median = .0094; interquartile range [IQR] = .0015) than between the Unfamiliar Masked and Unfamiliar Alone conditions (median = .0096; IQR = .0022) (Figure 2A; *S* = 88, *p* = .015, *Z* = 2.43).

**Figure 2.**
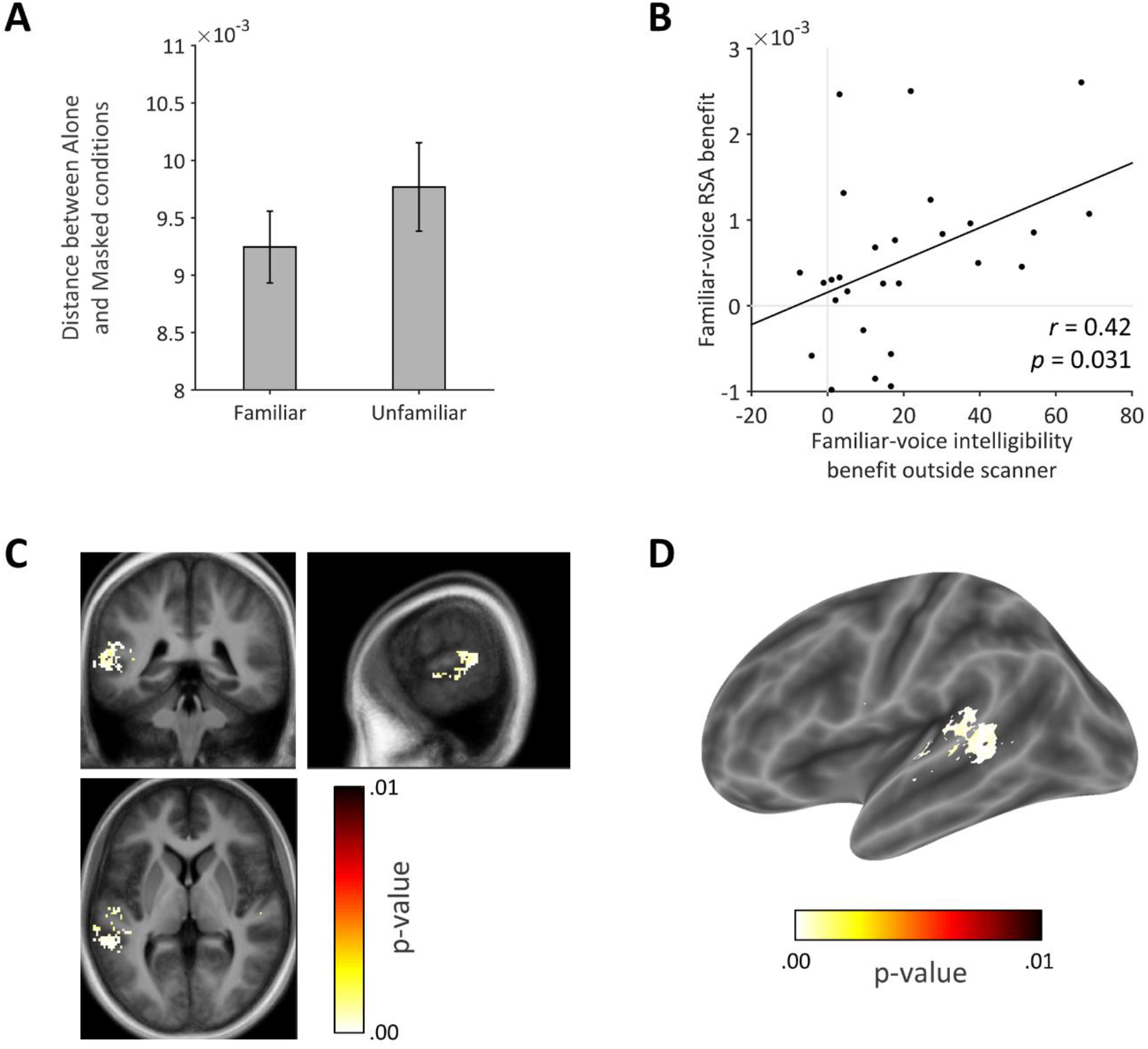
Functional MRI results (= 27). (A) Results from the Representational Similarity Analysis (RSA). The y-axis shows the correlation distance metric between the Alone and Masked conditions, plotted separately for conditions in which the target sentence was Familiar or Unfamiliar. (B) Correlation between the familiar-voice benefit to intelligibility (i.e., difference in percent correct between the Familiar Masked and Unfamiliar Masked conditions) measured in the post-scan behavioural task and the familiar-voice RSA benefit (i.e., difference in the RSA distance metric between the Familiar and Unfamiliar conditions). (C,D) Areas identified in searchlight RSA, displayed on sections from the average structural image (44 participants) and on an inflated cortical surface, plotted using BSPMVIEW (51).

### Robustness of neural activity correlates with intelligibility benefit

Figure 2B shows the relationship between the Familiar-Unfamiliar RSA difference and behavioural performance outside the scanner, across participants. The Familiar-Unfamiliar RSA difference was significantly correlated with the magnitude of the intelligibility benefit that participants gained from their familiar voice [*r* = .42, *p* = .031; 95% CI = .04–.69].

### Intelligibility pattern is most prominent in posterior STG, MTG and PT

We used searchlight RSA to find the brain areas that were most sensitive to the Familiar-Unfamiliar RSA difference. Figure 2C–D shows the results of this analysis, thresholded at *p* < .05 FWE. Significant voxels were present in left posterior STG, planum temporale (PT), and MTG.

To check if these significant areas overlapped with primary auditory cortex, we compared the maps in Figure 2C with left Te1.0 defined by Morosan et al. (23). Auditory cortex activity was posterior and/or inferior to area Te1.0, implying that significant effects of familiarity occur outside primary auditory cortex.

### No evidence for difference in regions for familiar vs unfamiliar voices

For completeness, we also analysed the data using a standard univariate approach, using a threshold of *p* < .05 FWE. No voxels were significant at this threshold for the main effect of Familiarity or for the interaction between Familiarity (Familiar or Unfamiliar) and Masker (Alone or Masked). However, this was probably not due to a lack of power, because we found a number of significant regions for the main effect of Masker, possibly reflecting differences in acoustics or effort. The peak voxels in these regions are listed in Table 1. The voxels showing greater activity for the Masked than Alone condition (i.e., two talkers > one talker) with the lowest p-values (*p*_*FWE*_ < .001) were located in PT, posterior supramarginal gyrus, angular gyrus, superior parietal lobule, middle frontal gyrus, inferior frontal gyrus, superior frontal gyrus, paracingulate gyrus, and the frontal operculum. Voxels showing greater activity for the Alone than Masked condition were located in the paracingulate gyrus, frontal pole, and left hippocampus.

**Table 1.** Results from the univariate contrast between the Alone and Masked conditions. Statistical analyses were conducted at the group level using one-sample t-tests, and were thresholded at p = .05 after correcting for family-wise error (FWE). Peak locations were labelled using the Harvard-Oxford atlas based on the MNI co-ordinates.

| Contrast | Peak location | t | $p_{FWE}$ | MNI co-ordinates (mm) | | |
| --- | --- | --- | --- | --- | --- | --- |
|  |  |  |  | x | y | z |
| Masked > Alone | Planum Temporale | -13.24 | < .001 | -55 | -21 | 4 |
|  | Supramarginal Gyrus (posterior) | -10.19 | < .001 | -55 | -42 | 11 |
|  | Middle Frontal Gyrus | -13.11 | < .001 | -42 | 17 | 32 |
|  | Middle Frontal Gyrus | -8.32 | < .001 | -40 | 3 | 54 |
|  | Inferior Frontal Gyrus, pars opercularis | -8.03 | .001 | -50 | 19 | 11 |
|  | Superior Parietal Lobule | -12.44 | < .001 | -31 | -57 | 46 |
|  | Planum Temporale | -10.21 | < .001 | 62 | -19 | 7 |
|  | Superior Frontal Gyrus | -10.13 | < .001 | -5 | 12 | 58 |
|  | Paracingulate Gyrus | -6.06 | .031 | 10 | 28 | 35 |
|  | Angular Gyrus | -9.63 | < .001 | 35 | -55 | 46 |
|  | Middle Frontal Gyrus | -9.21 | < .001 | 47 | 31 | 33 |
|  | Frontal Operculum Cortex | -8.18 | < .001 | 35 | 24 | 4 |
|  | Right Cerebral White Matter | -7.31 | .002 | 31 | 50 | 4 |
|  | Right Caudate | -7.07 | .004 | 12 | 10 | 7 |
|  | Location not in atlas | -6.95 | .005 | -38 | -62 | -33 |
|  | Left Cerebral White Matter | -6.36 | .017 | -29 | 42 | 4 |
|  | Left Caudate | -6.06 | .031 | -12 | 12 | 4 |
|  | Middle Frontal Gyrus | -5.97 | .037 | 35 | 10 | 63 |
|  | Left Cerebral White Matter | -5.96 | .038 | -12 | 7 | 4 |
|  | Right Cerebral White Matter | -5.95 | .039 | 24 | 54 | -5 |
|  | Left Cerebral White Matter | -5.84 | .049 | -14 | 3 | 4 |
| Alone > Masked | Paracingulate Gyrus | 8.97 | < .001 | -5 | 54 | 0 |
|  | Frontal Pole | 8.24 | < .001 | 0 | 59 | 23 |
|  | Left Hippocampus | 8.35 | < .001 | -29 | -31 | -12 |
|  | Temporal Pole | 8.05 | .001 | 52 | 7 | -30 |
|  | Supramarginal Gyrus (anterior) | 6.73 | .008 | 55 | -29 | 32 |
|  | Left Cerebral White Matter | 6.49 | .013 | -14 | -52 | 28 |
|  | Frontal Pole | 6.32 | .018 | -16 | 45 | 49 |
|  | Frontal Pole | 5.97 | .038 | 10 | 54 | 44 |
|  | Subcallosal Cortex | 5.95 | .039 | -3 | 7 | -9 |
|  | Frontal Pole | 5.91 | .042 | 9 | 55 | 40 |
|  | Frontal Pole | 5.91 | .042 | 14 | 52 | 47 |
|  | Subcallosal Cortex | 5.85 | .048 | -2 | 10 | -9 |

## Discussion

Our results demonstrate that multivariate brain activity evoked by speech is more resilient to competing speech when it is spoken by someone familiar compared to when the talker is unfamiliar. The extent to which familiar voices were associated with more robust multivariate brain activity co-varied with the magnitude of the behavioural intelligibility benefit that participants gained from their familiar voice. Thus, multivariate BOLD activity indexes the intelligibility benefit that people gain from a familiar voice in the presence of a competing talker. In this experiment, the unfamiliar talkers that we presented were familiar to other participants, so the familiar-voice advantage must be due to familiarity with a friend or partner’s voice, and probably cannot be explained by different voice acoustics. Our results are consistent with the idea that experience-driven changes in intelligibility are reflected in distributed activity in nonprimary auditory cortical areas.

The searchlight analysis showed that the difference between familiar and unfamiliar voices is most prominent in left posterior STG and MTG. On one hand, we might have expected the difference to be most prominent in higher-level areas, such as the inferior frontal cortex because this area has been implicated in effortful listening (11, e.g., 12, 24–26)—and speech spoken by familiar people is easier to understand, so may be associated with less effort. On the other hand, STG activity has been found in previous studies that have manipulated intelligibility by changing speech acoustics (11–13, 27) and lexical predictability (14). In addition, Gennari et al. (28) found that phoneme discrimination was associated with increased activity in posterior STG and MTG under conditions of lower than higher cognitive load. Therefore, the differences we found could reflect lower load associated with processing a voice that is familiar, compared to a voice that is unfamiliar. Indeed, behavioural studies are consistent with the idea that familiar voices use fewer cognitive resources to process (29, 30), which would have a greater effect in the presence of a competing talker than when familiar and unfamiliar voices are presented alone.

Bilateral STG, STS, and MTG have been shown to respond more to vocal than non-vocal sounds, and they have been previously labelled as ‘temporal voice areas’ (31–34). Our finding that left STG is sensitive to the difference between familiar and unfamiliar voices is consistent with this idea, and further suggests that the degree of familiarity with someone’s voice affects activity in these temporal voice areas. It is worth noting that all of our conditions contained voices, so our results do not reflect voice-specific processing, but rather activity that differs depending on how familiar a voice is to a particular individual. Furthermore, our results demonstrate that this is due to improved intelligibility of familiar voices rather than familiarity *per se*, because the RSA effect significantly correlated with the intelligibility benefit. We counterbalanced the voices in the familiar and unfamiliar conditions so that voice identities were similar across conditions. The area of STS we found was most sensitive to the familiar-unfamiliar voice difference is located more posteriorally than the temporal voice areas in some of the previous studies (e.g., 31–33). Nevertheless, Bethmann, Scheich, and Brechman (35) found that the greatest difference between familiar and unfamiliar voices occurred in both anterior and posterior areas that were more distant from primary auditory cortex, which is consistent with our results. In addition, Warren et al. (36) proposed that abstraction of voice identity information occurs in posterior STS, and subsequent analysis occurs in anterior parts of STS and MTG. Birkett et al. (37) found greater activity for familiar than unfamiliar voices in the left MTG. However, all of these studies either used tasks that asked participants to judge voice familiarity (35, 37) or had participants passively listen to stimuli while speaker identity varied across conditions (36)—whereas the task in the current study was to report the words that were spoken.

Based on previous neuropsychological studies showing that deficits to familiar voice recognition and unfamiliar voice discrimination arise from different patterns of brain damage (38–40), we might have expected to find that familiar and unfamiliar voices preferentially activated different regions of the brain. However, our univariate analysis showed no significant differences, which may have arisen because our task was based on discriminating the content of speech (i.e., the words that were spoken), rather than recognising the voice: This is consistent with behavioural evidence showing that people use familiar-voice information in different ways when the goal is to understand the words spoken by someone familiar than when the goal is to recognise someone’s identity from their voice (2), and neural evidence that brain activity differs depending on whether the task is one of intelligibility or voice recognition (41–43).

Our results are consistent with the idea that speech spoken by familiar and unfamiliar people is processed in similar regions of the brain, but speech spoken by familiar people is more resistant to background noise than speech spoken by familiar people. This could occur, for example, if the amplitude envelope of speech is encoded more precisely (44, 45) for familiar than unfamiliar voices at early stages of auditory processing, or if neuronal gain (46) increases for frequency channels corresponding to the frequencies of a familiar voice. Cognitively, this could be underpinned by better knowledge of acoustic properties of a familiar than unfamiliar voice—for example, due to better inferences of vocal tract properties associated with the formants in speech (e.g., 47) or from processes operating on the masker that are related to a reduction in informational masking (see 29, 45), which could potentially arise from lower cognitive load for processing familiar than unfamiliar voices.

An interesting question for future research is whether other (cognitive or acoustic) benefits to intelligibility are associated with similar patterns of brain activity. For example, the benefits from matching visual word primes (14), increases in TMR (e.g., 48, 49), or spatially separating target and competing speech (48, e.g., 50). Given that manipulating acoustic factors and visual word primes lead to intelligibility-related activity in broadly similar regions of the brain that we found to be maximally sensitive to the familiar-voice benefit to intelligibility, it is possible that similar mechanisms could underlie these intelligibility benefits (i.e., across different domains). On the other hand, acoustic and cognitive factors could affect intelligibility through different processes, which ultimately lead to similar activity reflecting the improvement in intelligibility via either mechanism. In this paper, we have demonstrated that the multivariate pattern of activity, analysed using RSA, is sensitive to differences in intelligibility between familiar and unfamiliar voices; this method may, therefore, be useful for comparing neural instantiations of the intelligibility benefits gained through different (cognitive or acoustic) means.

## Conclusions

Overall, the current study demonstrates that more intelligible speech is associated with multivariate activity in nonprimary auditory cortical areas. We found the greatest sensitivity to voice familiarity in posterior superior and middle temporal gyri. We found the magnitude of the BOLD difference in dissimilarity between familiar and unfamiliar voices covaries with the intelligibility benefit that participants gain from their friend or partner’s voice. Therefore, this paper identifies a neural substrate for the (cognitive) benefit to speech intelligibility that occurs when speech is spoken by someone familiar.

## Materials and Methods

### Participants

We recruited 27 participants (9 male, 22 right-handed), who had taken part in a previous behavioural experiment on voice familiarity, and who had a friend or partner who had been recorded speaking a list of sentences. Participants were 19–68 years old (median = 22 years, inter-quartile range = 6), were native English speakers, and had average pure-tone audiometric thresholds better than 20 dB HL in each ear (measured at four octave frequencies between 0.5 and 4 kHz). The experiment was cleared by Western University’s Health Sciences Research Ethics Board. Informed consent was obtained from all participants.

### Design

First, participants completed an adaptive behavioural task to determine the target-to-masker ratio (TMR) for reporting 40% of sentences correctly when both talkers were unfamiliar. During the subsequent scanning session, all stimuli were presented at the adapted TMR—which ensured that the intelligibility level of the baseline (unfamiliar) condition was equivalent for all participants.

During the scanning session, we presented 6 experimental conditions in a 3 × 2 factorial design. Target sentences were either spoken by a familiar (“Familiar”) or by one of two unfamiliar (“Unfam-1” and “Unfam-2”) talkers. The unfamiliar talkers in the scanning session were different than those presented in the pre-scan behavioural task, to prevent participants becoming overly familiar with particular unfamiliar voices. During the scanning session, target sentences were either presented alone (“Alone”) or in the presence of a competing sentence (“Masked”). Masking talkers were always unfamiliar and different from the target talker. In addition, we included silent trials that contained no acoustic stimuli.

Finally, we conducted a post-scan behavioural task to measure the intelligibility of the materials heard in the three Masked conditions in the scanner, which provided an independent measure of the familiar-voice benefit to intelligibility for each participant. Sentences from the three conditions (Familiar Masked; Unfam-1 Masked; Unfam-2 Masked) were presented in a randomized order.

### Apparatus

The pre- and post-scan behavioural sessions were conducted in a quiet room. Acoustic stimuli were presented through a Steinberg Media Technologies UR22 sound card and were delivered binaurally through Grado Labs SR225 headphones. Participants viewed visual stimuli on the monitor of a Lenovo ThinkPad P50 20EN laptop and responded using a mouse.

While participants were in the MRI scanner, acoustic stimuli were presented through the same Steinberg Media Technologies UR22 sound card, which was connected to a stereo amplifier (PYLE PRO PCA1 for 22 participants, PYLE PRO PCAU22 for 5 participants). Acoustic stimuli were delivered binaurally through Sensimetrics insert earphones (Model S14 for 22 participants, Model S15 for 5 participants) and were presented at a comfortable listening level that was the same for all participants. Visual stimuli were projected onto a screen at one end of the magnet bore, which participants viewed through a mirror attached to the head coil.

### Stimuli

Acoustic stimuli were spoken sentences that had been recorded by each participants’ friend or spouse in a previous experiment. Sentences were from the Boston University Gerald (BUG) corpus (19), which follow the structure: “<Name> <verb> <number> <adjective> <noun>”. In the sub-set of sentences used in the experiment, there were two names (‘Bob’ and ‘Pat’), eight verbs, eight numbers, eight adjectives, and eight nouns, which are displayed in Figure 1A. An example is “Bob brought three red flowers”.

Sentences were recorded using a Sennheiser e845-S microphone connected to a Steinberg Media Technologies UR22 sound card. The recordings were conducted in a single-walled sound-attenuating booth (Eckel Industries of Canada, Ltd.; Model CL-13 LP MR). The sentences had an average duration of 2.5 seconds (*s* = 0.3). The levels of the digital recordings of the sentences were normalised to the same root mean square (RMS) power.

During the experiment, each participant heard sentences spoken by their familiar partner and sentences spoken by eight unfamiliar talkers, who were the partners of other participants in the experiment. Sentences spoken by six of the unfamiliar talkers were presented in the pre-scan behavioural adaptive test, and sentences spoken by the other two unfamiliar talkers were presented in the scanning session and post-scan behavioural test.

We planned to present each voice to one participant (i.e. their partner) as a familiar talker and to two other participants as an unfamiliar talker. However, this was not possible because the partners of 8 people did not participate in this experiment. Thus, 8 voices were presented as unfamiliar but never as familiar, 10 voices were presented only once as familiar and once as unfamiliar, and 3 voices were only presented as familiar. In total, we used 36 different talkers. Thus, across the group, familiar and unfamiliar conditions were acoustically similar.

### Procedure

#### Pre-scan behavioural

To determine the target-to-masker ratio (TMR) for reporting 40% (chance = 0.02%) of sentences correctly, we used a weighted up-down procedure (Kaernbach, 1991). On each trial, participants heard two sentences from the BUG matrix spoken simultaneously by two different unfamiliar talkers of the same sex. They identified the four remaining words of the sentence that began with a particular target name (“Bob” or “Pat”), by clicking buttons on a screen (Figure 1A). The words in the masker sentence were always different to the words in the target sentence. We adapted the TMR in 3 separate, but interleaved, runs—which each contained a different pair of unfamiliar talkers. Each run stopped after 12 reversals and we calculated thresholds for each run as the median of the last 5 reversals. For each participant, we calculated the median of the thresholds across the three runs: this TMR value was used during the MRI session.

#### Functional MRI

During the MRI session, we presented 12 functional runs, each containing 25 trials (300 trials total). We presented 48 trials in each of the six experimental conditions, as well as 12 silent trials. All 7 trial types were pseudorandomly interleaved.

In three of the conditions, participants heard 48 sentences from the BUG matrix (19), which were either spoken by their familiar (“Familiar Alone”) or by one of their two unfamiliar (“Unfam-1 Alone” and “Unfam-2 Alone”) talkers. In the other three conditions (“Familiar Masked”, “Unfam-1 Masked” and “Unfam-2 Masked”), participants heard the same sentences spoken by the same three talkers, but they were presented simultaneously with a different sentence from the BUG matrix that was spoken by one of the two unfamiliar talkers. The onsets of the target and masker sentences were identical. For the two conditions in which one of the unfamiliar talkers was presented as the target, the masker sentence was spoken by the other unfamiliar talker. In the Familiar Masked condition, the Unfam-1 and Unfam-2 talkers were each presented as the masker talker on half of the trials. The words in the masker sentence were always different from those in the target sentence.

Figure 1B illustrates the trial structure: We modified the task so it was more amenable to responses inside the MRI scanner. On each trial, the target sentence was the one that began with a particular name word (‘Bob’ or ‘Pat’). Half of target sentences began with ‘Bob’ and the other half began with ‘Pat’. Acoustic stimuli were positioned such that the middle of the target sentence occurred 4 seconds before the beginning of the first volume collection of a pair of volumes (see Section 2.6, MRI data acquisition, below). At the beginning of each trial, the target name word was displayed visually on the screen (even when the target sentence was presented alone). The name word was presented on the screen for 300 ms at the beginning of each trial, then a fixation cross was presented for 3700 ms. Four seconds after the trial began, participants saw a probe sentence written on the screen. They were asked to indicate whether the probe sentence was the same as the target sentence they heard spoken. They held a button box in one hand and pressed one button if the probe sentence was the same and a different button if the probe sentence was different. The name word in the probe sentence was always the same as the target name. On half of trials, the other four words were also the same. On the other half of trials, one of the four words was different. On Alone trials, the different word was selected randomly from the other words in the BUG corpus. On Masked trials, the different word was from the masker sentence. The placement of the incorrect word in the sentence (i.e., 2^nd^, 3^rd^, 4^th^, or 5^th^ word) was counterbalanced across trials.

For the 12 silent trials, the visual cue word was “Rest”, and no acoustic stimuli were presented.

Immediately before the scanning session, participants completed a practice, which contained 14 trials with the same (fixed) TMR that was used in the MRI session. The practice was conducted in a quiet room with the same equipment as the pre-scan adaptive task. The trial structure was identical to the functional runs of the scanning session. Participants responded using two keys on the laptop.

#### Post-scan behavioural

Finally, participants completed a behavioural task outside the scanner. We presented three conditions in which there was always a competing masker: Familiar Masked, Unfam-1 Masked, and Unfam-2 Masked. The trials were identical to those presented in the MRI session, but they were presented in a different (pseudorandomly interleaved) order. The post-scan behavioural was divided into two halves: In one half, target sentences began with the name word ‘Bob’, and in the other, target sentences began with the name word ‘Pat’. The order of the name words was counterbalanced across participants. The structure of each trial was identical to the pre-scan adaptive part: participants identified the four remaining words from the target sentence by clicking buttons on a screen (Figure 1A). Participants completed 144 trials (48 in each of the three conditions), with a short break every 24 trials.

### MRI data acquisition

MRI was conducted on a 7.0 Tesla Siemens MAGNETOM scanner at Robarts Research Institute, Western University (London, Ontario, Canada) with a 32-channel receive coil. At the beginning of the session, we acquired a whole-brain T1-weighted anatomical image for each participant with the following parameters: MP2RAGE; voxel size = 0.75 mm isotropic; 208 slices; PAT GRAPPA of factor 3; anterior-to-posterior phase encoding, time-to-repeat (TR) = 6000 ms, echo time (TE) = 2.83 ms. T2*-weighted functional images were acquired using echo-planar imaging (EPI), with: voxel size = 1.75 mm isotropic; 63 slices; multi-band acceleration of factor 3 with interleaved slices; field of view of 208 mm; TR = 1000 ms; echo spacing = 0.45 ms; PAT GRAPPA of factor 3; posterior-to-anterior phase encoding; bandwidth = 2778 Hz/Px. Acquisition was transverse oblique, angled away from the eyes, and in most cases covered the whole brain. (If the brain was too large for the field of view, slice positioning excluded the very top of the superior parietal lobule.) We used interleaved silent steady state (ISSS) imaging (Schwarzbauer et al., 2006): Each trial contained 7 ‘silent’ scans (radio frequency pulses without volume acquisition) followed by 2 scans with volume acquisition (Figure 1B). Acoustic stimuli were presented during the silent period between volume acquisitions. We collected 52 volumes from each participant (2 per trial) in each of the 12 runs. The first two ‘dummy’ scans were presented immediately prior to the first trial of each run and were excluded from the analyses. We collected field maps immediately after the functional runs (short TE = 4.08 ms, long TE = 5.1 ms).

### Analyses

For the analyses, we collapsed across the conditions in which unfamiliar voices were presented as targets (i.e., “Unfam-1 Alone” and “Unfam-2 Alone”; “Unfam-1 Masked” and “Unfam-2 Masked”).

#### Behavioural data

We calculated sensitivity (d’) for target recognition performance during the MR session using loglinear correction (20), and chance d’ of 0.3. For the post-scan behavioural, we calculated the percentage of sentences in which participants reported all four words (after the name word) correctly. The data met the assumptions for normality, as assessed by non-significant Shapiro-Wilk and Kolmogorov-Smirnov tests, and by visual inspection of box plots and Q-Q plots. We used Pearson’s product moment correlation coefficients to compare d’ in the MRI session with percent correct in the post-scan behavioural session.

#### MRI data

MRI data were preprocessed using SPM12 (Wellcome Centre for Human Neuroimaging, London, UK). Each participant’s functional images (EPIs) were unwarped using their field maps and were realigned to the first image of the run. The functional and anatomical images were coregistered to the mean EPI, then normalised to the standard SPM12 template (avg305T1). For RSA analyses, we took the mean of the two adjacent volumes for each trial. For the univariate analyses, we took the same average after applying spatial smoothing, to ensure the data met the assumptions of Gaussian random field theory for multiple comparisons correction (21). For spatial smoothing, we used a Gaussian kernel with a full-width at half-maximum of 12 mm. We analysed the results from each participant at the first level using a General Linear Model with the motion realignment parameters and a regressor corresponding to each run as covariates of no interest.

For RSA, we entered the unsmoothed images into the first level analysis and extracted the betas from each participant that corresponded to the experimental conditions. The region of interest (ROI) was defined using the Neurosynth database: We used a meta-analysis of all studies (N = 81) that included the term ‘Speech Perception’ and used this to mask the imaging data. We analysed the ‘distance’ between the betas for pairs of conditions using MATLAB 2017b. We focussed on conditions in which the same sentences were spoken by the same talker, but in the presence or absence of a competing masker; for example, “Familiar Alone” compared with “Familiar Masked”. We performed the analyses once using correlations as the distance metric and once using Euclidean distances, and we obtained the same pattern of results using both methods. We, therefore, report results using correlation distances, which were defined as 1 minus the Pearson’s correlation coefficient. At the group level, we compared distances for Familiar and Unfamiliar conditions using signed-rank tests.

For each participant, we extracted the distances between the alone and masked stimuli in the Familiar and Unfamiliar conditions and used the difference as an index of the Familiar-Unfamiliar RSA difference. We then used a Pearson’s correlation, across participants, to examine the relationship between the Familiar-Unfamiliar RSA difference and the behavioural benefit to intelligibility that each participant obtained from their familiar voice. We calculated this behavioural benefit from the post-scan behavioural test, as the difference between percent correct in the Familiar Masked condition and the Unfamiliar Masked conditions.

We used the RSA toolbox (22) for the searchlight RSA analysis. We searched within the ‘Speech Perception’ Neurosynth mask for areas that were particularly sensitive to the difference in distances between Familiar and Unfamiliar conditions. We defined an expected dissimilarity matrix that contained a smaller value (0.5) for the Familiar Alone with Familiar Masked cells, than for the Unfamiliar Alone with Unfamiliar Masked cells (1.0). The remaining cells in the matrix were of no interest for this analysis and were, therefore, set to NaNs. We used the correlation distance metric on the betas at the individual subject level. To compare the expected dissimilarity matrix with the data, we used Spearman’s correlations. Our searchlight radius was 15 mm.

For the univariate analyses, we entered the spatially smoothed images into the first level analyses, where we applied our contrasts of interest: the main effect of Familiarity (Familiar or Unfamiliar), the main effect of Masker (Alone or Masked), and the interactions. We analysed the resulting contrast images at the group level using one-sample t-tests. All contrasts were corrected for family-wise error (FWE; 21).

## Acknowledgements

This work was supported by funding from the Canadian Institutes of Health Research (CIHR; Operating Grant: MOP 133450), the Natural Sciences and Engineering Research Council of Canada (NSERC; Discovery Grant: 327429-2012), and Western University’s Canada First Research Excellence Fund BrainsCAN initiative. We thank Joe Gati for his help piloting scanning parameters.

